# Swimming kinematics of rainbow trout behind cylinder arrays: the effect of vortex street periodicity and turbulence kinetic energy

**DOI:** 10.1101/2024.04.15.589564

**Authors:** David M. Sparks, Edwin Rajeev, Alberto Canestrelli, James C. Liao

## Abstract

Fish in the wild often contend with complex flows that are produced by natural and artificial structures. Research into fish interactions with turbulence often investigates metrics such as turbulence kinetic energy (TKE) or fish positional location, with less attention paid to the specific interactions between vortex organization and body swimming kinematics. Here we compare the swimming kinematics of rainbow trout (*Oncorhynchus mykiss*) holding station in flows produced by two different 3 x 5 cylinder arrays. We systematically utilized computational fluid dynamics to generate one array that produced a Kármán vortex street with high vortex periodicity and TKE (KVS array), and another that produced low periodicity and TKE, similar to a parallel vortex street (PVS array). The only difference in swimming kinematics between cylinder arrays was an increased tail beat amplitude in the KVS array. In both cylinder arrays, the tail beat frequency decreased and snout amplitude increased compared with the freestream.

The center of mass amplitude was greater in the PVS array than in only the freestream, however, suggesting some buffeting of the body by the fluid. Notably, we did not observe Kármán gaiting in the KVS array as in previous studies. We hypothesize that this is because (1) vorticity was dissipated in the region where fish held station in this study and (2) cylinder arrays produced vortices that were in-line rather than staggered. These results are the first to quantify the kinematics and behavior of fishes swimming in the wake of multiple cylinder arrays, which has important implications for biomechanics, fluid dynamics, and fisheries management.

**SUMMARY STATEMENT:** The swimming kinematics of rainbow trout are largely preserved across two, 3 x 5 cylinder array treatments that differed in vortex periodicity and turbulence kinetic energy.

## INTRODUCTION

Fishes living in currents commonly hold station in the complex flows found behind both natural objects such as rocks, corals, and submerged vegetation, and also man-made structures such as bridge pilings and dams. The cost of swimming in these flows varies: some studies show that fish exhibit higher oxygen consumption and decreased stability in turbulence (Enders et al., 2003; Tritico and Cotel, 2010; Webb, 1998). Other studies show that fish that hold station around a bluff body consume less oxygen compared to swimming in uniform flow (Liao et al., 2003a; Liao et al., 2003b; Przybilla et al., 2010; Taguchi and Liao, 2011). Kármán gaiting fish, for example, exploit the staggered, alternating vortices of a vortex street by slaloming between them (Liao, 2004; Liao et al., 2003b). Entraining and bow waking fish exploit local high-pressure regions around a cylinder to perform minimal swimming movements (Liao et al., 2003a; Przybilla et al., 2010).

Despite the prevalence of assemblages of bluff bodies in the current swept-environments inhabited by fishes, very little is known about the effect of their downstream wake on the mechanics of fish swimming. Yet extending our understanding of environmental energy recapture into more physically-complex habitats promises to provide new insights and strategies for efficient fish locomotion (Liao, 2022) and inform the design of fishway passages to minimize the impact on native ecosystems (Castro-Santos et al., 2009; Puzdrowska and Heese, 2019; Lacey et al., 2012; Wilkes et al., 2017). A fuller understanding of how fishes can benefit energetically from swimming in these complex habitats will require a discrete examination of the details of the flow structure that fish depend on, such as vortex size or periodicity (Liao, 2003a,b; Enders et al., 2003; Stewart et al., 2016).

A great diversity of wakes can be produced behind multiple cylinders based on the ratio of streamwise to cross-stream cylinder gap space (Gao et al., 2020). We ran Computational Fluid Dynamics (CFD) simulations on 500 cylinder arrays of differing spacing ratios to search the parameter space of wakes based on vortex shedding periodicity (Stewart et al., 2016). From this search, we selected two different wakes from two different arrays: the first array exhibited a discreet Kármán vortex street, and consequently high vortex periodicity and turbulence kinetic energy (TKE), and is herein called the KVS array. The second array exhibited low vortex shedding periodicity and TKE, resembling a parallel vortex street (Karasudani and Funakoshi, 1994), and is herein called the PVS array. These two cylinder arrays were then fabricated and assembled in a laboratory flume according to those CFD simulations to examine the effect of their wakes on the swimming kinematics of rainbow trout.

This study employs a combination of systematically selected CFD simulations with live experiments on rainbow trout to directly test the effects of specific wake parameters on station holding swimming kinematics.

## MATERIALS AND METHODS

### Overview

Rainbow trout were exposed to two cylinder arrays, as well as a control treatment with no cylinders (freestream flow). Each of the three treatments were run at three flow speeds (22 cm s^-1^, 48 cm s^-1^, and 74 cm s^-1^) in a 175 L recirculating flow tank (Loligo Systems, Tjele, Denmark). Computational fluid dynamics (CFD) simulations for one cylinder array predicted a highly periodic flow with a discreet Kármán vortex street, called the KVS array (Fig. 1A). Simulations for the other cylinder array predicted a low periodic flow, which included similar features to a parallel vortex street, called the PVS array (Fig. 1B). A high-speed camera captured images of the ventral silhouette of fish at 100 fps (Phantom Miro 340, Vision Research, Wayne, NJ, USA). A machine learning program (DeepLabCut) was then trained on 260 annotated frames at 600,000 iterations to recreate the outline of the fish in each frame. Using customized Matlab scripts, fish midlines were then reconstructed from these outlines and used to calculate body kinematics such as tail beat frequency, body wavelength, snout amplitude, center of mass (COM) amplitude, and tail tip amplitude (Liao et al., 2003a).

**Figure 1.**
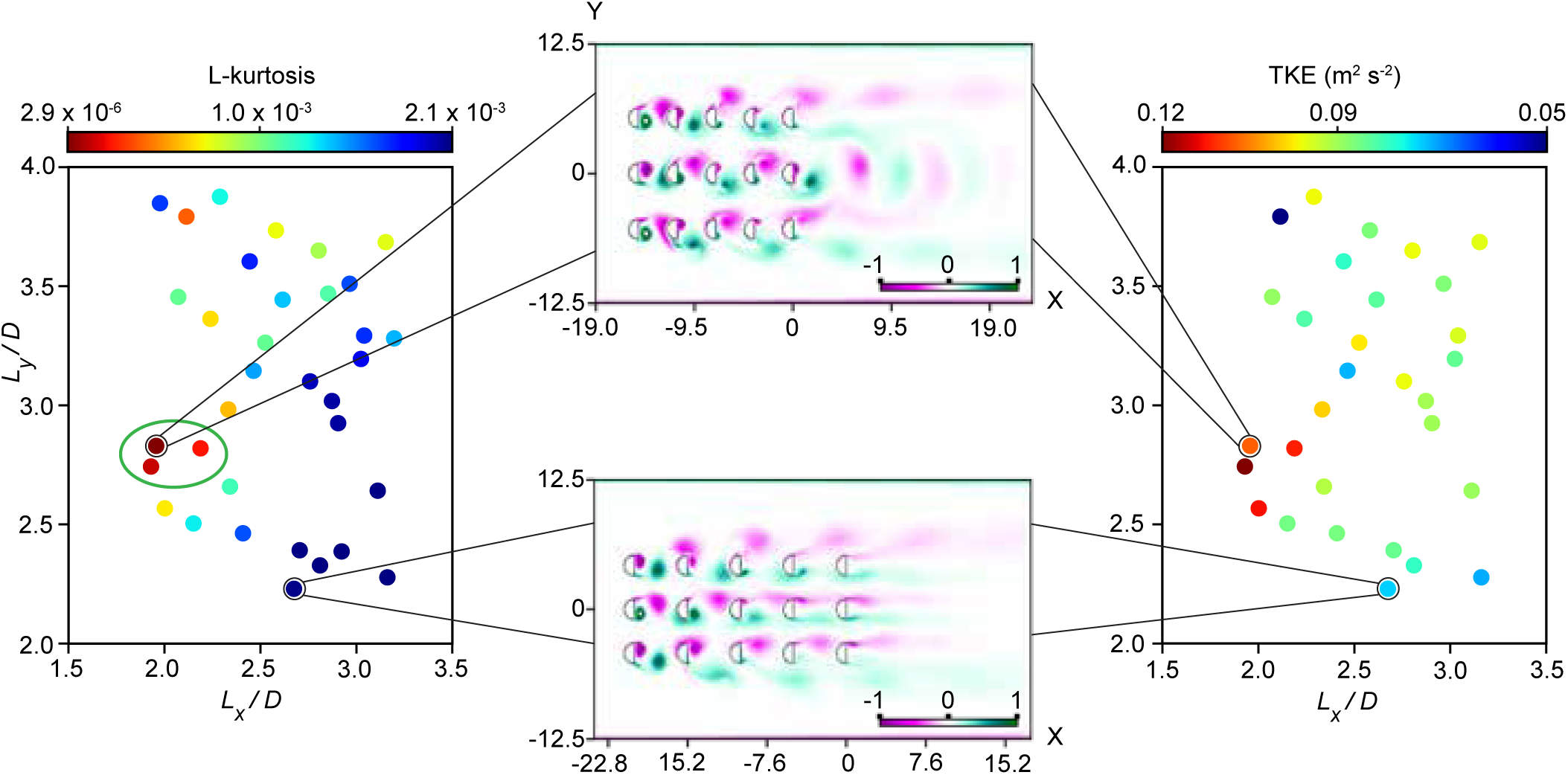
Optimization study results and turbulence kinetic energy of CFD simulations. (Left) The values of the objective function (L-kurtosis of the FFT spectrum) and (right) turbulence kinetic energy are plotted against the spacing ratio (*L_x_/D* and *L_y_/D*). Cylinder arrays with the most optimal spacing ratios are encircled in green. (Middle top) Optimal (KVS array) and (middle bottom) sub-optimal (PVS array) vorticity contours selected for the experiment. Color gradient scale represents vorticity (s^-1^). The experimental cylinder array was constructed based on these two arrangements. (Y and X axes in centimeters).

### Trout acquisition and husbandry

Thirty juvenile rainbow trout (*Oncorhynchus mykiss*) were obtained from Wolf Creek National Fish Hatchery, KY, USA and held in a 500 L recirculating round tank maintained at 15 ± 1 °C using an in-line chiller (Delta Star chiller, model DS-4-TXV, Aqua Logic, San Diego, CA, USA). A continuous flow was created to accustom animals to swimming against current. Trout were allowed to acclimate to this environment for a minimum of 48 hours before experimentation. Five individual rainbow trout were randomly selected for experimental trials (mean body length ± s.e.m. = 7.4 ± 0.1 cm)

After data were collected for the first cylinder array, the fish was placed into a separate holding tank for the remainder of the study to ensure that it could be identified for subsequent experiments. Holding chambers were constructed out of 30 cm long PVC piping with netting, suspended in the main holding tank with other fish, and fish were fed a maintenance diet. This step was taken to avoid moving the cylinder arrays mounted in the flow tank, which could alter the hydrodynamics of the wake by disrupting their precise configuration.

### Hydrodynamic treatment selection

Two cylinder arrays were selected based on the wakes predicted across 120 simulations: (1) a highly periodic flow with a discreet Kármán vortex street (KVS array; Fig.1A) and (2) a low periodic flow with similar features to a parallel vortex street (PVS array; Fig. 1B).

#### Computational fluid dynamics

Two-dimensional flow fields were modeled around cylinder arrays using computational fluid dynamics (CFD) in OpenFOAM (v2012). The CFD model numerically solved the Navier-Stokes equations that govern incompressible fluid motion and a 2-D Shear Stress Transport (SST) k-ω Unsteady Reynolds Averaged Navier-Stokes (URANS) turbulence model was adopted.

A finite volume method with an overset grid solved the incompressible Navier-Stokes equations, which accounts for maintaining constant momentum. The walls of the half-cylinders were discretized using a structured grid and divided into 130 grid points. Due to the complex nature of vortex interaction between cylinders inside an array, turbulence was fully resolved at the cylinder wall. The effects of the tank side walls were also taken into account for the cylinder-vortex interactions to closely mimic laboratory conditions. No-penetration and no-slip boundary conditions were prescribed at the cylinder surfaces and the side walls. A velocity-inlet boundary condition was prescribed upstream of the cylinders, and a free boundary condition downstream. The inlet velocity was prescribed at a nominal flow speed equivalent to 74 cm s^-1^ (Reynolds number = 14,300).

The CFD model was validated by comparing the Strouhal number (St) and the instantaneous wake wavelengths of vortex streets with the experimental results obtained by Stewart et al. (2016). The St was obtained from the dominant frequency of a Fast Fourier Transform (FFT) of the vertical component of the velocity vector (Uy) with respect to time. The instantaneous wake wavelength was determined by the spatial wavelength of Uy along the cylinder axis extending downstream of the cylinders. The instantaneous wake wavelength was computed at each time step using the FFT of Uy with respect to the x−axis behind the cylinder. A grid convergence study was also conducted to ensure that numerical results would be independent of mesh size, time step, and the computational domain.

The average vortex shedding frequency for each of the two selected arrays at 74 cm s^-1^ was based on the St and physical parameters of the experimental tank by the following equation:

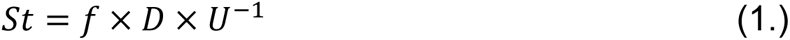

Whereby *f* was solved for in terms of *D*, individual cylinder diameter, and *U*, the flow velocity in the region of the cylinders. To account for solid blocking effects causing flow constriction near the cylinders, *U* was calculated by:

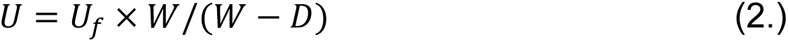

Where *W* is the width of the tank, and *U_f_* is the nominal flow speed.

#### Optimization

An optimization procedure was performed to find the cylinder arrangement that maximizes Kármán vortex street periodicity because of the prevalence of Kármán gaiting in highly periodic wakes (Akanyeti and Liao, 2013a; Stewart et al., 2016).

Sixty simulations were run to determine the optimal values of the design variables. Due to the large number of design variables and the non-linear nature of the dynamics, we sought an efficient optimization method. We employed Surrogate Based Optimization (SBO) to approximate the objective function in the entire parameter space starting from 55 numerical simulations, called sample points (Adams et al., 2020). In this case, the objective function was the L-kurtosis of the FFT spectrum peakedness, which is periodicity.

In our optimization procedure, an initial set of random sample values of the design variables were first created to describe the design space. The design variables for this specific problem are the spacing parameters, *L_x_/D* and *L_y_/D*, which are the ratio of cylinder gap space in the streamwise and in the cross-stream directions, respectively, to cylinder diameter. Following this, evaluations of the objective function on each sample point were performed in OpenFOAM. These evaluations were subsequently used to build the surrogate model upon which the optimal value could quickly be computed via an interpolation method. DAKOTA (Adams et al., 2020) was used to provide an interface between CFD (OpenFOAM) and analysis methods (SBO).

### Experimental setup

#### Experimental area

Fish swimming experiments took place in a 175 L recirculating flow tank (Loligo Systems, Tjele, Denmark) with a working flume area of 30 cm height x 25 cm width x 89 cm length maintained at 15 ± 1 °C (Stewart et el., 2016). A high-speed camera (100 fps at 1440 x 2560 pixel resolution, Miro LAB340, Vision Research, Wayne, NJ, USA) was aimed at a 45° mirror to image the ventral view of the flow tank. An LED light source (Lyra DMX, IkanCorp) placed above a white plexiglass sheet was used as a diffuser to create a high-contrast silhouette of the fish.

#### Treatment conditions

The frame of the multiple-cylinder arrays was modeled in SolidWorks (version 2020) (Fig. 2, A). The design was then manufactured out of clear acrylic to minimize visual stimuli for fish (Liao, 2006) by the UF Infinity Fab Lab (College of Design, Construction, and Production, University of Florida, Gainesville, FL, USA). The frame allowed cylinders to be independently adjusted relative to each other, such that they could be disassembled and reconstructed to produce new arrangements. The arrays were constructed within ± 1 mm of the CFD simulations (Fig. 2, B). Assembled arrays were lowered into the flow tank and secured to resist lift from the flow.

**Figure 2.**
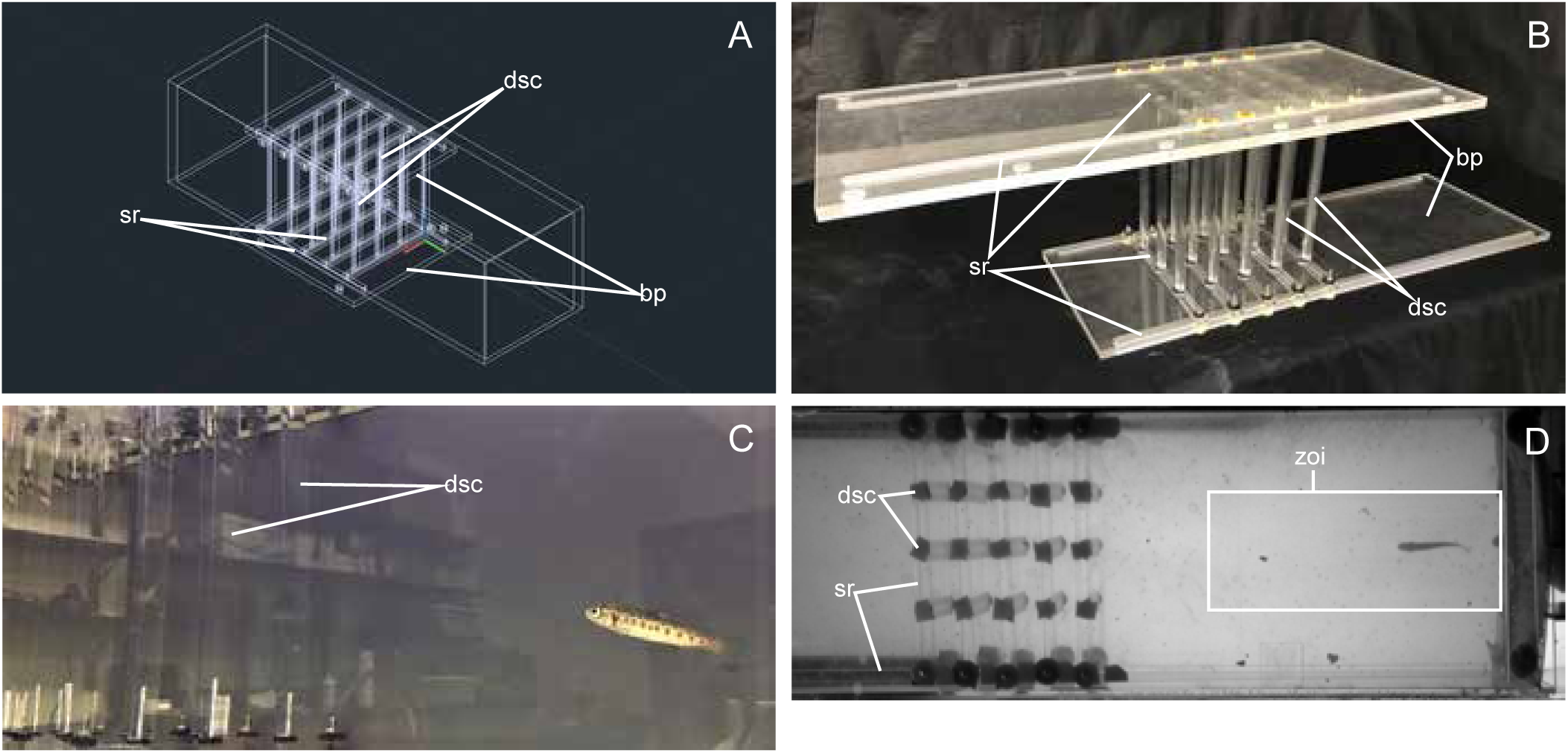
Experimental set up. (A) CAD drawings of parts for an adjustable array of multiple D-shaped cylinders were designed using SolidWorks. (B) Parts were laser cut out of clear acrylic by UF Infinity Fab Lab and cylinders were purchased separately. Arrays were manually constructed to replicate both CFD simulation cylinder arrays within ± 1 mm. (C) Five rainbow trout (7.4 ± 0.1 cm) (*Oncorhynchus mykiss*) each individually swam downstream from each combination of hydrodynamic treatment (freestream, Kármán vortex street, parallel vortex street) and flow speed (22 cm s^-1^, 48 cm s^-1^, and 74 cm s^-1^). (D) High-speed video recordings of 16 tail beats of the ventral silhouette were taken for each individual x hydrodynamic treatment x flow speed combination within a pre-determined zone of interest. (Key: dsc = D-shaped clinders, sr = sliding rails, bp = baseplates, zoi = zone of interest).

Rainbow trout were exposed to two cylinder arrays and a control treatment of freestream flow (no cylinder array) at each of three flow speeds (22 cm s^-1^, 48 cm s^-1^, and 74 cm s^-1^). The lowest velocity represents the flow speed at which trout were introduced to the flow tank. The highest flow speed was experimentally determined to be the fastest rate at which fish would hold station. The intermediate flow speed was selected for a third point of comparison.

### Experimental procedure

Individual trout were transported from the holding tank and introduced downstream to the cylinder array with the flow tank set at the lowest velocity treatment (Fig. 2C, D). A fixed, rectangular “zone of interest” was determined from the development of vortices in vorticity contours generated by CFD simulations models. This zone occupied a certain downstream distance from the last column of cylinders and was used to guide the collection of video sequences for subsequent analysis of swimming kinematics (Fig. 2D). If a swimming fish did not hold station and drifted back against the downstream baffle, it was returned to its holding chamber to rest and experiments were continued at a later time. Sixteen total tail beats were collected from each individual for each flow speed and hydrodynamic treatment.

Upon completion of the experiment, trout were euthanized by an overdose of MS-222 and its total length measured. The anteroposterior center of mass (COM) was also determined by iteratively balancing the fish between two probes and measuring the balance point (Liao et al., 2003a).

### Digitization

The body silhouette of each fish was identified using DeepLabCut (version 2.1.10.1) (Mathis et al., 2018; Nath et al., 2019) with ipython (version 7.20.0) (Fig. 3). Training data were developed from 260 frames in which the snout, tail tip, and six points on the left and right side of the fish outline were annotated in varying luminance and noise (i.e. bubbles). The data were trained for 600,000 iterations before predicting locations on the remaining data. Predicted locations were graphically and visually inspected for gross anomalies using a custom script in R (version 4.0.4). Data were then manually grouped into sequences of four tail beats in R. A midline was then reconstructed, and body kinematics calculated with a custom MatLab script (version R2020a). If >10% of data from a video file were inaccurate, the outline of the fish was manually digitized with a separate custom MatLab script.

**Figure 3.**
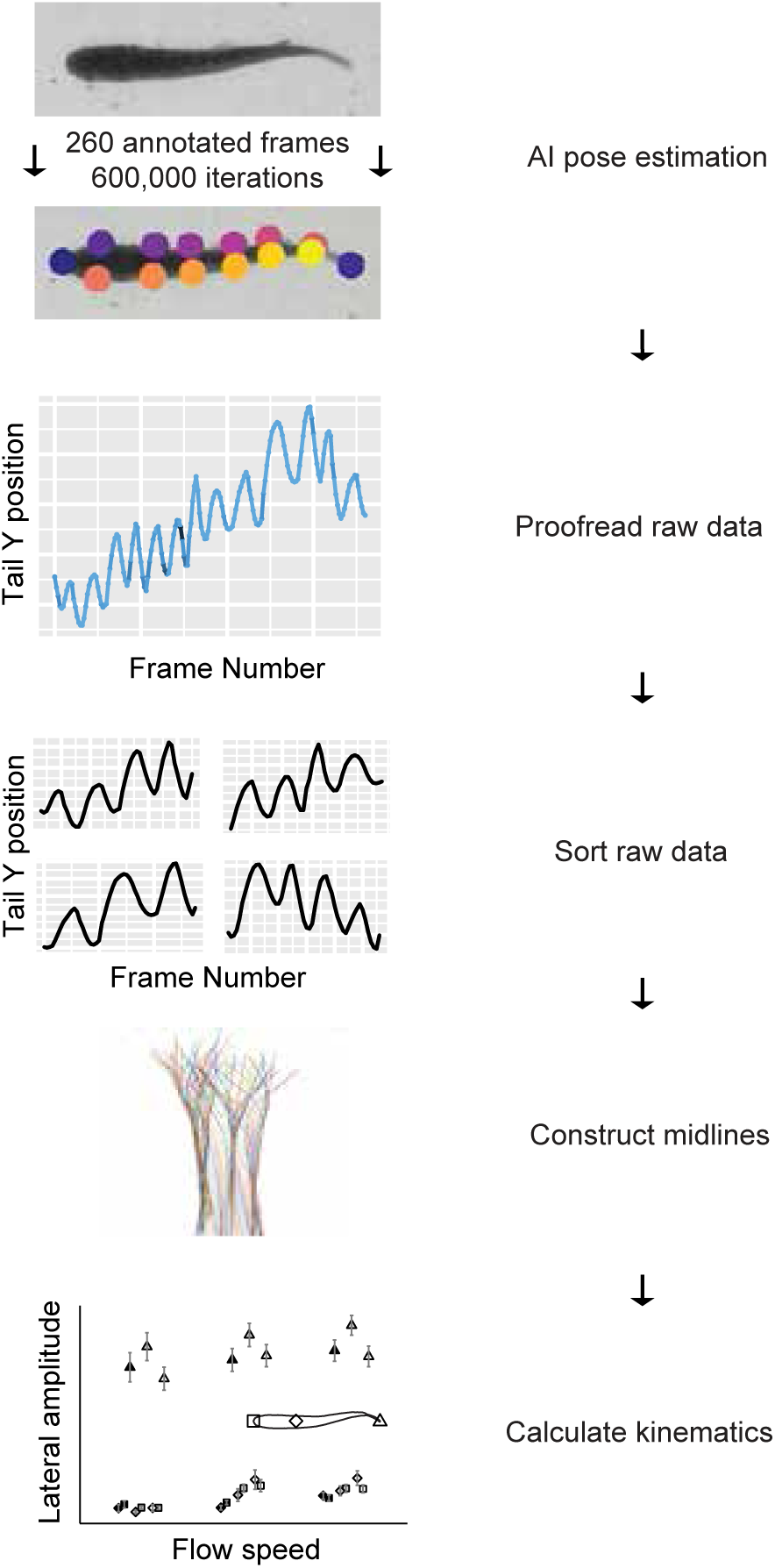
Schematic concept of the computational workflow. Annotations of the snout, tail tip and 6 points along both sides of the fish’s body were placed on 260 representative frames in DeepLabCut (ipython). A library was made by training DeepLabCut on those frames for 600,000 iterations. DeepLabCut then estimated the annotated body positions on all video data. These estimations were manually proofread for extreme, obvious miscalculations by DeepLabCut, then sorted into sets of four tail beats (R). The midlines of the fish were constructed, and thence kinematics were calculated (MatLab).

Tail beat frequency, body wavelength, and the amplitude at three locations along the body (snout, COM, and tail tip) were calculated using MatLab. Tail beat frequency was determined from the inverse of the average period (time between maxima and between minima of tail beats). Wavelength was calculated as the average phase speed (determined from the mean speed of maxima traveling along the midline) divided by the tail beat frequency. All amplitudes were calculated by halving the distance between maxima and minima along the midline (i.e. maximum lateral excursion from the midline).

### Statistical analysis

Amplitude data were log transformed to satisfy normality. One datum was an outlier and removed from the snout amplitude data because it was clear from reviewing the video that the fish was not holding station, and one set of four tail beats was removed for the same reason. All statistical analyses were conducted in R (Kassambara, 2021). We calculated means of kinematics with s.e.m. (Table 1), then ran two-way ANOVAs with an FDR adjustment to compare hydrodynamic treatments and flow speeds. Where significance was found, we conducted Tukey’s HSD post-hoc tests (Tukey-Kramer HSD in the case of the snout amplitude) at 95% confidence to determine differences by either main effect or interaction effect. We further conducted one sample, left-tailed t-tests to compare the mean tail beat frequencies of both cylinder arrangements at 74 cm s^-1^ with their respective vortex shedding frequencies.

**Table 1.**
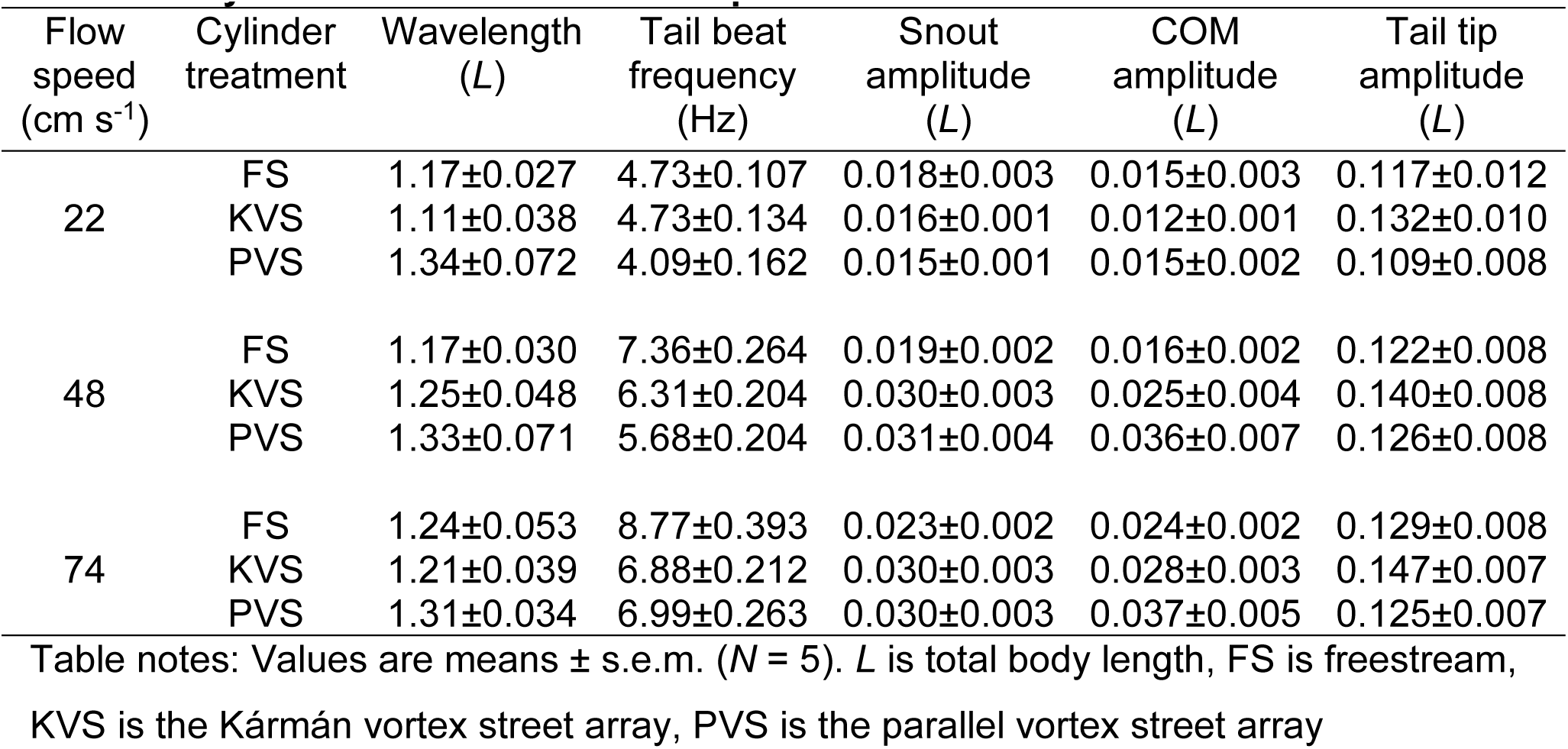
Summary statistics of trout kinematic variables for each combination of cylinder treatment and flow speed.

| Flow speed<br>(cm s<sup>-1</sup>) | Cylinder treatment | Wavelength<br>(L) | Tail beat frequency<br>(Hz) | Snout amplitude<br>(L) | COM amplitude<br>(L) | Tail tip amplitude<br>(L) |
| --- | --- | --- | --- | --- | --- | --- |
| 22 | FS | 1.17±0.027 | 4.73±0.107 | 0.018±0.003 | 0.015±0.003 | 0.117±0.012 |
|  | KVS | 1.11±0.038 | 4.73±0.134 | 0.016±0.001 | 0.012±0.001 | 0.132±0.010 |
|  | PVS | 1.34±0.072 | 4.09±0.162 | 0.015±0.001 | 0.015±0.002 | 0.109±0.008 |
| 48 | FS | 1.17±0.030 | 7.36±0.264 | 0.019±0.002 | 0.016±0.002 | 0.122±0.008 |
|  | KVS | 1.25±0.048 | 6.31±0.204 | 0.030±0.003 | 0.025±0.004 | 0.140±0.008 |
|  | PVS | 1.33±0.071 | 5.68±0.204 | 0.031±0.004 | 0.036±0.007 | 0.126±0.008 |
| 74 | FS | 1.24±0.053 | 8.77±0.393 | 0.023±0.002 | 0.024±0.002 | 0.129±0.008 |
|  | KVS | 1.21±0.039 | 6.88±0.212 | 0.030±0.003 | 0.028±0.003 | 0.147±0.007 |
|  | PVS | 1.31±0.034 | 6.99±0.263 | 0.030±0.003 | 0.037±0.005 | 0.125±0.007 |

Table notes: Values are means ± s.e.m. ( $N = 5$ ). $L$ is total body length, FS is freestream, KVS is the Kármán vortex street array, PVS is the parallel vortex street array

## RESULTS

### CFD and optimization

The goal of the CFD simulations was to identify cylinder arrangements that optimally generate periodic Kármán vortex wakes within and downstream of a 3 x 5 cylinder array. The objective function, L-kurtosis, was maximized within this optimization, which quantifies the periodicity of Kármán vortex streets. This analysis gives an improved understanding of the influence of the spacing-ratio in the streamwise (*L_x_/D*) and cross-stream (*L_y_/D*) directions on Kármán vortex street formation.

The interplay between the streamwise and cross-stream spacing ratios was responsible for achieving optimal Kármán vortex streets, such that there was no single optimal spacing ratio. Instead, a distribution within the parameter space conveyed a range of values for *L_x_/D* (1.9-2.5) and *L_y_/D* (2.4-3) that promotes the formation of optimized Kármán vortex streets (Fig. 1C).

In this optimal range, vortical structures generated between cylinders held their shape as they translated downstream. The subsequent Kármán vortex street formation downstream of the array also preserved its shape with high periodicity. We selected a Kármán vortex street from this range with a Strouhal number of 0.21 and a periodicity of 0.1059 m^2^ s^-1^ (L-kurtosis: 0.0063) (KVS, Fig. 1A). Outside of this range, the large vortical structures generated from the most upstream cylinders failed to maintain their shape as they translated downstream. This phenomenon is likely due to the improper impingement of the vortical structures on the downstream cylinders, thus causing a disrupted flow field downstream of the array. Hence, vortex street formation downstream of the arrangements that were outside of the optimal range were lacking in periodicity. We selected a flow field with similar features to a parallel vortex street (Karasudani and Funakoshi, 1994) in which vortices were separated cross-stream from the center cylinder’s streamwise axis. The Strouhal number was 0.23 and the periodicity was 0.00557 m^2^ s^-1^ (L-kurtosis: 5.39 x 10^-5^) (PVS, Fig. 1B). Periodicity for both arrays were greater than 3 x 10^-5^ m^2^ s^-1^, the maximum value considered by a previous study looking at fish swimming behind two cylinders (Stewart et al. 2016).

After experiments were conducted, we calculated the turbulence kinetic energy (TKE) of all CFD simulations directly with OpenFOAM’s built-in program (Fig. 1D). The TKE of the KVS and PVS array was 0.10453 m^2^ s^-2^ and 0.07076 m^2^ s^-2^, respectively.

### Swimming kinematics across hydrodynamic treatments and flow speeds

Neither flow velocity (F = 3.02, p = 0.0514) or the interaction term with flow treatment (F = 0.676, p = 0.6098) had a significant effect on body wavelength after FDR adjustment (α = 0.0333, Fig. 4). Flow treatment did have a main effect on wavelength (F = 3.54, p = 0.0312), although no significance was found in the post-hoc test. Conversely, there was a significant interaction effect between flow treatment and flow velocity on tail beat frequency (F = 4.527, p = 0.00168). Both flow velocity and flow treatment had main effects on snout amplitude (F = 24.972, p = 3.1 x 10^-10^; F = 4.166, p = 0.0171, respectively) and center of mass (COM) amplitude (F = 23.003, p = 1.42 x 10^-9^; F = 4.495, p = 0.0125, respectively), though the effect of flow treatment was smaller. Flow velocity (F = 3.987, p = 0.0203) and flow treatment (F = 5.79, p = 0.00368) also had an effect on tail tip amplitude, but flow treatment had a greater effect than flow velocity, which was the opposite pattern of the anterior body amplitudes. The interaction effects on snout (F = 1.995, p = 0.0975), COM (F = 1.682, p = 0.1564), and tail tip (F = 0.124, p = 0.97358) amplitudes were not significant.

**Figure 4.**
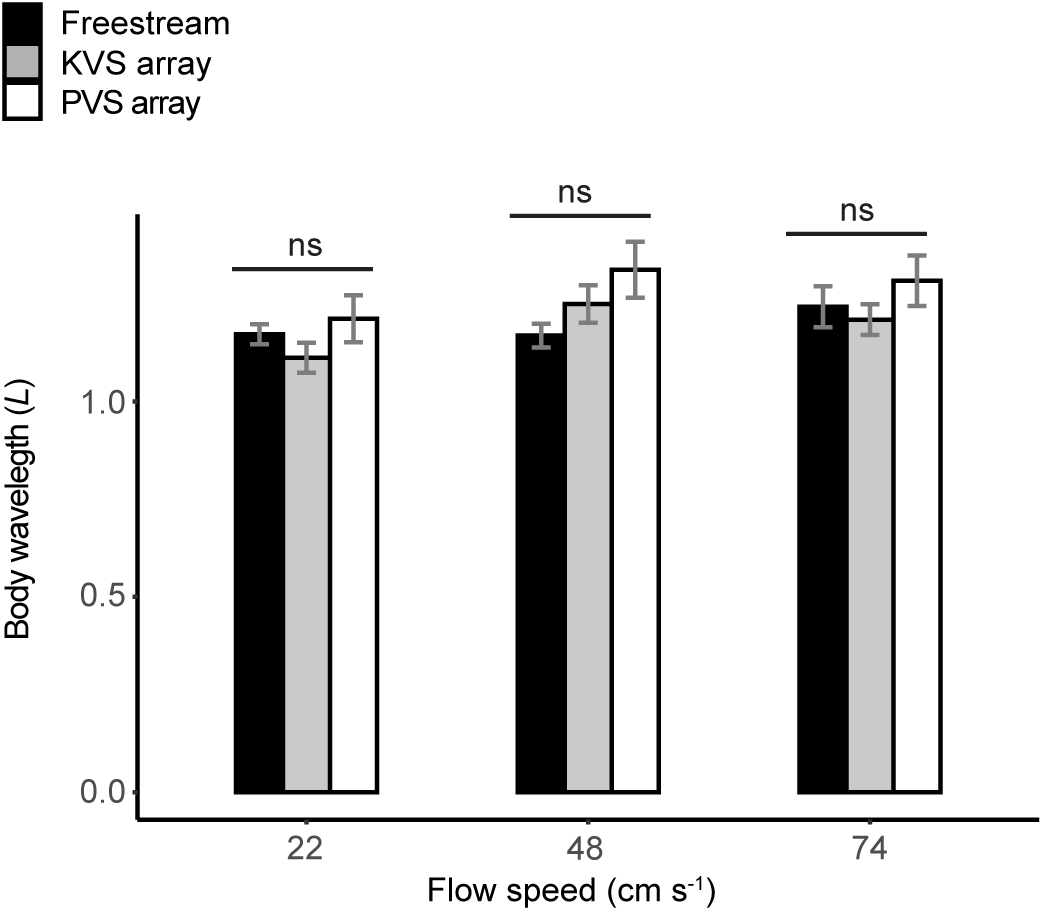
Mean body wavelength (*L* = fish body length) and s.e.m. bars (95% error) across flow speeds and cylinder treatments. Shade represents hydrodynamic treatment (black = freestream; grey = Kármán vortex street; white = parallel vortex street). An FDR-adjusted two-way ANOVA detected a main effect on wavelength due to hydrodynamic treatment (reported in body of results), but no significance was found by a Tukey’s post-hoc test.

Tail beat frequency generally increased with flow velocity, though was generally lower behind cylinder arrangements than in the freestream (Tukey’s test: α = 0.05, Fig. 5). Within flow treatments, there were differences between each velocity and either other velocity, except within the KVS array between 48 cm s^-1^ and 74 cm s^-1^. There were no significant differences between flow treatments at 22 cm s^-1^. At 48 cm s^-1^, the tail beat frequency in the freestream was significantly higher than that in the PVS array (p = 2.48 x 10^-5^) and the KVS array (p = 0.0398) but did not differ significantly between the PVS and KVS arrays. Similarly at 74 cm s^-1^, the tail beat frequency was significantly higher in the freestream compared with both the PVS (p = 7.08 x 10^-6^) and the KVS arrays (p = 1.66 x 10^-6^), but did not significantly differ between the PVS and KVS arrays. This points to an increasing tail beat frequency with velocity, but an overall lower rate of increase when trout swam downstream to the multiple cylinder array compared with the freestream.

**Figure 5.**
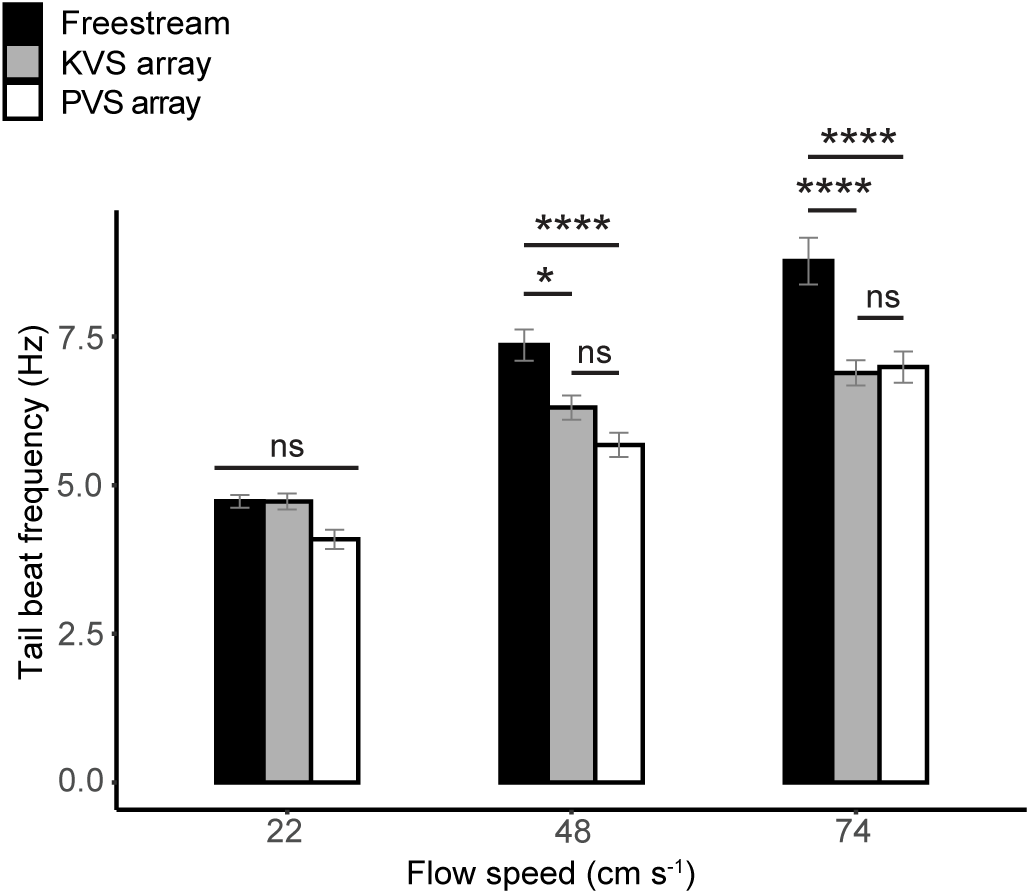
Mean tail beat frequency (Hz) with s.e.m. bars (95% error) across flow speeds and cylinder treatments. Shade represents hydrodynamic treatment (black = freestream; grey = Kármán vortex street; white = parallel vortex street). Tukey’s adjusted significance levels according to Tukey’s post hoc test following a FDR-adjusted two-way ANOVA are represented by asterisks (p < 0.0001 = ****; p < 0.05 = *; ns = not significant). Also notable, frequency differed significantly (p < 0.0001) between each flow speed and either other flow speed within each hydrodynamic treatment, except for between 48 cm s^-1^ and 74 cm s^-1^ in the Kármán vortex street.

Tail tip amplitude differed most significantly between the PVS array and the KVS array (p = 0.00618). There was also a significantly larger tail tip amplitude in the KVS array than in the freestream (p = 0.018), but the amplitude did not significantly differ between in the PVS array and the freestream. Among velocities, there was one significant difference in tail tip amplitude between the lowest and the highest flow speeds (p = 0.0202), and no other significant differences. Overall, it seemed that the KVS array had the greatest effect on the tail tip amplitude.

The most significant increases in COM and snout amplitudes were between the lowest flow speed and the two highest speeds (i.e. maximum p-value = 9.20 x 10^-5^ between 22 cm s^-1^ and 48 cm s^-1^ for the COM) (Fig. 6). The COM amplitude also differed significantly between the two highest speeds (p = 0.0438), but the snout amplitude did not. Among hydrodynamic treatments, the COM amplitude only differed significantly between the freestream and the PVS array (p = 0.00886). The snout amplitude, however, differed significantly between the freestream and both the PVS (p = 0.0434) and the KVS arrays (p = 0.0296). Though the interaction terms of the ANOVAs were non-significant, it is qualitatively apparent from Figure 6 that variation between hydrodynamic treatments was more pronounced at the two highest speeds than the slowest speed. Minimally, both velocity and the cylinder arrangements affected amplitudes on the anterior body.

**Figure 6.**
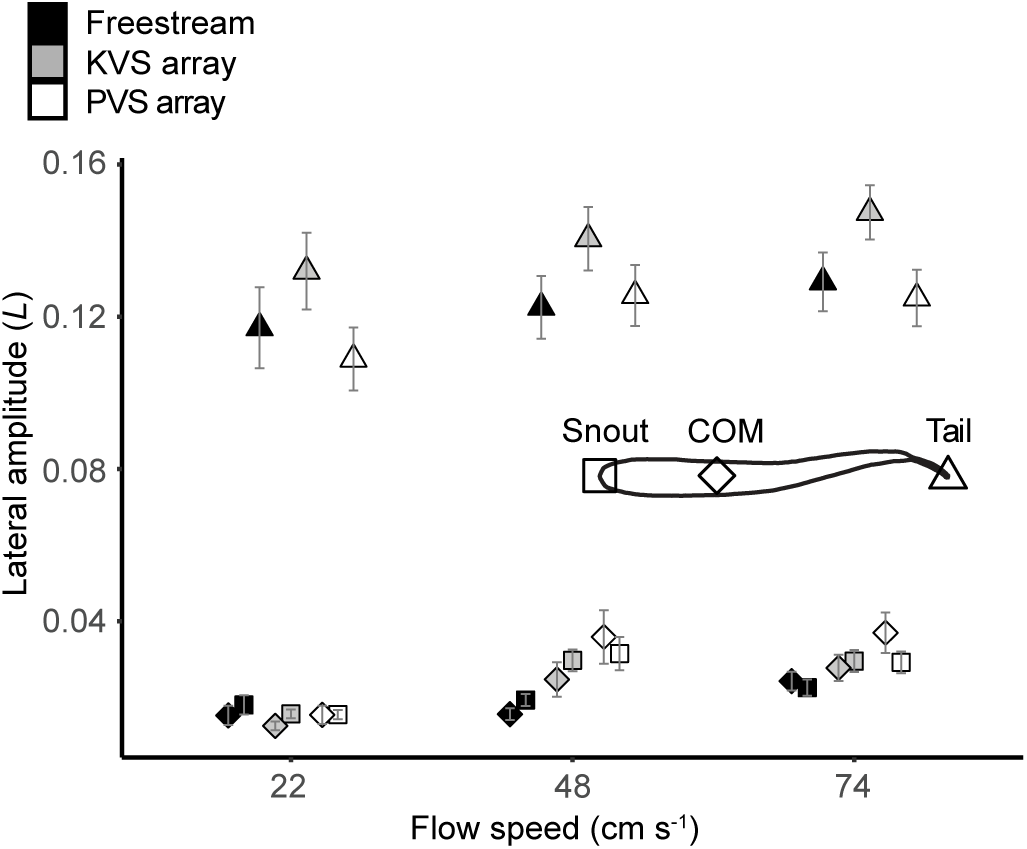
Mean lateral amplitude (*L* = fish body length) with s.e.m. bars (95% error) across flow speeds and cylinders treatments for three different locations along the body. Shade represents hydrodynamic treatment (black = freestream; grey = Kármán vortex street; white = parallel vortex street) and shape represents body location (square = snout; diamond = center of mass [COM]; triangle = tail tip). Results of a FDR-adjusted two-way ANOVA with a Tukey’s post hoc test are reported in the body of the chapter.

The vortex shedding frequencies of the KVS and PVS arrays at 74 cm s^-1^ were 8.85 Hz and 9.69 Hz, respectively. The tail beat frequencies in both cylinder arrangements were significantly lower than their respective vortex shedding frequencies (KVS array: p = 8.931 x 10^-9^; PVS array: p = 1.679 x 10^-9^).

### Qualitative observations of swimming behaviors

Though we set out to analyze station-holding behaviors, we observed other behaviors such as forward acceleration, entraining, wall following, and vertical movements through the water column. Among these behaviors were lateral trajectories we refer to as “casting”. When we increased flow velocity, trout made successive sweeping lateral excursions, sometimes spanning the entire flume (Fig. 7). This was followed by swimming directly to the downstream edge of the cylinder array, where trout would momentarily entrain behind a single cylinder before accelerating through the entire cylinder array (Fig. 8). Once upstream of the cylinder array, it swam with similar motions to bow waking, yet unlike bow waking, held station upstream of the gap between cylinders rather than directly in front of a cylinder.

**Figure 7.**
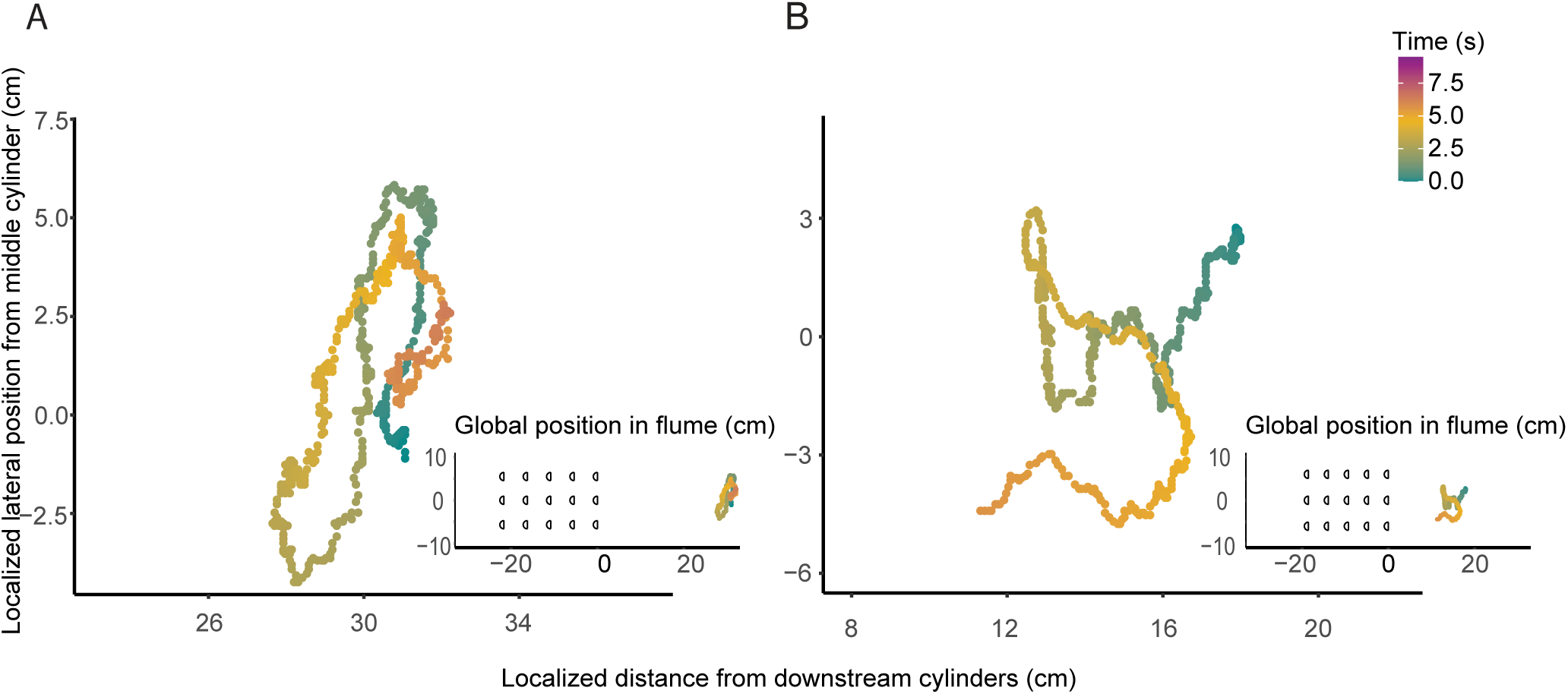
Cases of casting behavior in rainbow trout illustrated by digitized snout position across time in seconds (color). All X-Y axis units are 1:1. These instances are at a flow speed of 48 cm s^-1^ in the (A) parallel vortex street and (B) Kármán vortex street. Corresponding global position to the multiple cylinder array is shown by inset graphs.

**Figure 8.**
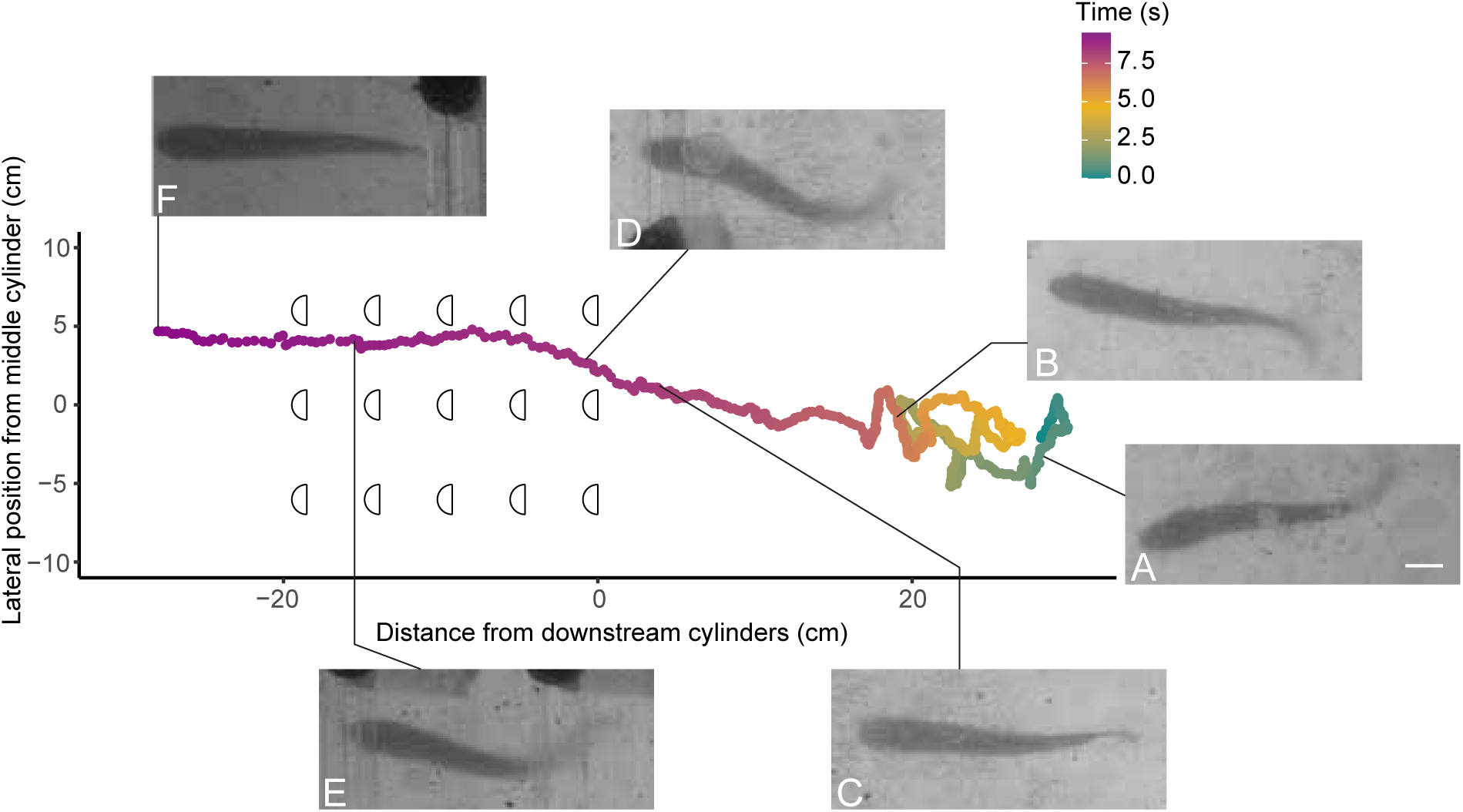
A representative, continuous recording of a casting event followed by acceleration through the cylinder array for a single fish in a Kármán vortex street at 74 cm s^-1^. X-Y axis units are 1:1 and time scale is the same as in Fig. 7. Image panels (A – F) are zoomed-in video frames of the trout swimming at specific time points indicated by the lines that connect the graph to the panels. The 1 cm scale bar in (A) is consistent across all panels. (A, B) Casting behavior, in which the trout angles its body against the flow and swims left or right in the flume. (C) Following casting, the trout swims quickly and directly to the middle downstream cylinder. The trout lingers and appears to entrain behind the cylinder edge for a moment. (D) Trout accelerates into the cylinder array and (E) continues to swim through the array with the same swimming movement, in this case briefly pausing twice. (F) Once past the cylinder array, trout behaves similarly to panel (C), though the position would imply bow waking. (Observed out of camera view) Trout resumes a plethora of behaviors shortly after (F).

## DISCUSSION

### Hydrodynamics behind cylinder arrays

CFD simulations revealed novel flow patterns behind multiple cylinders arranged in a 3 x 5 pattern. Unlike a single cylinder where the wake contains vortices that have a similar diameter to the cylinder, our results support recent analyses which show that it is possible for vortices in the wake of multiple cylinders to be larger in diameter than any single cylinder (Gao et al., 2020). In the KVS array of this study, the drag wake behind the central cylinder draws in vortices from lateral cylinders. These vortices circulate in the same direction as the vortices behind the central cylinder, resulting in elliptical vortices that display an increase in diameter in the cross-stream direction. Holding the streamwise cylinder spacing constant and increasing the cross-stream spacing results in the opposite effect, instead producing dissipated vorticity behind the central cylinder and coherent lateral vortex streets.

The range of cylinder spacings which were optimal for periodic Kármán vortex streets also possessed higher turbulence kinetic energy (TKE) than most other cylinder spacings (Fig. 1, Right), likely due to the presence of coherent vortices. In the PVS array, the flow field is characterized by dissipated vortices and the TKE is correspondingly lower.

### Swimming kinematics of trout behind cylinder arrays

#### Body undulation while holding station in a vortex street and freestream

Swimming trout hold station in steady flows by passing a mechanical body wave from head to tail with increasing lateral amplitude (Bainbridge, 1962; Di Santo et al., 2021; Webb, 1988). In the presence of a coherent Kármán vortex street, these kinematics change significantly, and fish adopt a Kármán gait (Liao et al., 2003a; Liao et al., 2003b). This mode of locomotion occurs when the length of the fish is between two to four times the diameter of the shed cylinder vortices. The Kármán gait is identifiable by large lateral amplitudes across the body, a longer body wavelength, and a decreased tail beat frequency that is slightly higher than the vortex shedding frequency (Liao et al. 2003a, Akanyeti and Liao 2013a, Stewart et al. 2016). Kármán gaiting fish slalom between vortices as they exploit each vortex’s energy in a largely passive way, with little to no axial muscle activity (Liao, 2007; Liao et al., 2003b; Taguchi 2010). A model of Kármán gaiting reveals that it is freestream swimming kinematics superimposed onto a vortex street, which generates greater lateral body translations, body rotations, and head yaw motions (Akanyeti and Liao, 2013b).

We expected that fish would Kármán gait in the KVS array, but their station holding kinematics did not meet the criteria for Kármán gaiting (Akanyeti and Liao, 2013b). Kármán gaiting fish synchronize their tail beat frequency with the cylinder vortex shedding frequency, but in this study the tail beat frequency was lower than the expected vortex shedding frequency behind both cylinder arrangements (Akanyeti and Liao, 2013a; Liao et al., 2003a). The body wavelength did not significantly differ between any treatments, including the KVS array and freestream (Fig. 4). The snout, COM, and tail tip amplitudes were also lower than that of Kármán gaiting fish (less than 80%, 80%, and 50%, respectively; Akanyeti and Liao, 2013a; Liao et al., 2003a). We suggest that the absence of large lateral translations of the body indicates that fish are swimming through, rather than slaloming between, cylinder vortices. Overall, the kinematics of trout holding station behind our cylinder arrays are similar to freestream swimming, to the point where swimming movements could be considered modifications of freestream swimming rather than movements resembling Kármán gaiting.

One possible reason for the absence of Kármán gaiting is that vortices were not as coherent as expected in the zone of interest where we collected kinematics data. Maintaining trout in one location for long enough periods of time to record their swimming motions was often challenging since trout commonly casted (Fig. 7). Additionally, if a trout was close enough to the cylinder array, it typically accelerated through the array (Fig. 8), which is why the zone of interest was distanced from the downstream cylinders. Since vortex strength diminishes with distance from a bluff body (Tritico and Cotel, 2010), the strength of the wake may have diminished in the zone of interest to a level where the trout would not Kármán gait.

Another explanation involves the organization and morphology of vorticity contours. A fish in a Kármán vortex street ‘slaloms’ between vortices, where there is adequate space between vortices in both the streamwise and cross-stream directions for fish to swim. The streamwise spacing of vortices in the KVS array of this study was comparable to that of other studies in which fish Kármán gait (Liao et al., 2003a; Liao et al., 2003b; Stewart et al., 2016). In the cross-stream direction, however, there was no space between vortices because of their oblong morphology and exceptional cross-stream overlap (Fig. 1, Middle-top).

#### Comparison of swimming kinematics across cylinder arrays

Despite potentially large differences in wake periodicity and TKE, only tail amplitude increased significantly in the KVS array compared to the PVS array and freestream treatments. This pattern is similar to Kármán gaiting fish in a vortex street, though the amplitudes in the Kármán gait are relatively larger (Akanyeti and Liao, 2013a; Akanyeti and Liao, 2013b; Liao et al., 2003a; Stewart et al., 2016). In the KVS array, vortices are in-line and TKE values are high, thereby creating a condition in which fish can intercept stronger, successive vortices of alternating circulation. The PVS array, in contrast, has low TKE and vortices that separate in the cross-stream direction; effectively, it establishes a condition where the x-axis behind the central cylinder does not experience the main effects of what is already relatively weaker vorticity.

It is unclear whether the higher tail amplitude in the KVS array is due to passive buffeting or actively controlled by muscular activity. Tail muscles have been shown to be inactive in Kármán gaiting (Liao, 2004), and even a dead trout will exhibit an increased tail amplitude (Beal et al., 2006), so it is possible that the tail is buffeted passively in the KVS array. However, the fishes’ overall swimming kinematics in this study best resemble a fish swimming in a freestream, which requires sequential muscle activity (Altringham and Ellerby, 1999; Gillis, 1998; Hammond et al., 1998; Jayne and Lauder, 1995; Liao 2004; Wardle et al., 1995). Furthermore, a dead trout (Beal et al., 2006) towed in a Kármán vortex street passively synchronizes its tail beat frequency to the vortex shedding frequency, whereas the tail beat frequency of the present study was significantly less than the vortex shedding frequency. This indicates a level of muscular control, whether to increase the tail amplitude or to prevent the tail amplitude from further increasing. Both mechanisms could also offer interesting implications for vortex exploitation given that work on both passively oscillating foils (Wu and Chwang, 1975), and harmonically oscillating foils (Triantafyllou et al., 2000) have shown the capacity for vorticity control and thrust production in a vortex street. Regardless, the proximity of strong, successive vortices in the KVS array appears to play an important role on tail beat amplitude and indicate some level of muscular control that is not present in the PVS array.

The COM amplitude in a swimming fish increases with speed (Xiong and Lauder, 2014), and can play an important role in stability (Lauder and Drucker, 2004; Webb and Weihs, 2015). We found that the COM amplitude was also greater in the PVS array than in the freestream, yet did not significantly differ in either hydrodynamic treatment from the KVS array. These amplitudes may reflect the lateral translation of the body through the interaction with vortices, rather than increases in body wave amplitude during undulation since there were no differences in wavelength among hydrodynamic treatments. Translational movements of the COM likely have little impact on swimming stability (Webb, 2002), while large lateral translations can help to exploit vortex recapture during Kármán gaiting (Liao, 2004; Liao et al., 2003a).

The lower periodicity of vortical flows behind tandem cylinders are less predictable to fish and can be punctuated by more chaotic flow (Stewart et al., 2016; Igarashi, 19681). Given the low TKE of the PVS array, it is likely that these chaotic flows accumulate below the integral scale, since the majority of vortices in turbulence are small (Tritico and Cotel, 2010). This combination of low periodicity and low TKE in the PVS array therefore produces a subtle and less predictable flow, which may affect control of the COM. If correct, it could explain the kinematic difference seen in the PVS array from the freestream, even though the COM amplitude in the KVS array does not differ from either the PVS array or the freestream.

Compared to swimming in the freestream, tail beat frequency decreased and snout amplitude increased when cylinders were present, yet did not differ between cylinder arrays. Tail beat frequency was also lower than the expected vortex shedding frequency in both cylinder arrays. This suggests that trout did not synchronize their tail beats to vortices as has been observed when Kármán gaiting (Akanyeti and Liao, 2013b; Liao et al., 2003a). To investigate whether tail beat frequency was simply the result of swimming in the reduced velocity behind the cylinders, we calculated the tail beat frequency of a fish swimming in the freestream at the value of the reduced flow produced by each cylinder array. We did not observe a difference in the tail beat frequency between the reduced flow speed and the experimental array, indicating that the decreased tail beat frequency may be due to reduced flow. This would not be the case for the increased snout amplitude, however, because snout amplitude decreases with flow speed (Table 1). It is more likely, then, that variables of the wake itself affected the snout amplitude. While the mechanism is unclear in our experiment, lateral motions of the snout can produce thrust when brook trout (*Salvelinus fontinalis*) swim in a freestream (Lucas et al., 2020). It would be interesting to investigate the interactions of the anterior body with different unsteady flows to explore alternative methods of thrust production in cylinder wakes.

We expected that swimming kinematics behind cylinder arrays would differ from freestream swimming, but we did not expect that they would differ in similar ways between arrays because of the substantial differences in periodicity, TKE, and vorticity. The underlying mechanisms remain unclear, but it is notable that (1) tail beat amplitude is the only kinematic which differed between cylinder arrays, (2) COM amplitude only differed between the PVS array and freestream, and (3) snout amplitude and tail beat frequency changed in opposite directions (increase vs. decrease) in response to the presence of cylinders. Flow visualization and electromyography would provide valuable insight in future studies (Liao, 2004; Liao et al., 2003a; Liao et al., 2003b; Stewart et al., 2016; Tritico and Cotel, 2010).

### Casting behavior

We commonly observed sweeping behaviors whereby previously station-holding trout swam in the cross-stream direction when exposed to higher flow velocities behind cylinder arrays (Fig. 7). These sweeping movements spanned tens of centimeters and were often followed by upstream acceleration through the cylinder array (Fig. 8). We hypothesize that sweeping allows trout to explore the source and bounds of their fluvial environment, similar to how moths cast to sample pheromone plumes (Kennedy, 1983). Wilks et al. (2017) found that Atlantic salmon move quickly around their experimental arena to sample the flow. Cross-stream sweeping has also been described in Mexican tetras (Elder and Coombs, 2015) and Giant Danio (Bak-Coleman et al., 2013) in a variety of flow and sensory (lateral line, visual) conditions. Although this study focused on station-holding kinematics, the motivation of fishes in turbulence may be to escape upstream. Casting is a robust behavior that should be considered in future studies looking at the ability of fishes to navigate unsteady flows behind complex physical structures.

### Summary

We selected two cylinder arrays that exhibited flows of differing periodicity, TKE, and vortex street organization using CFD modeling. We then fabricated these cylinder arrays and experimentally placed live fish behind them. Fish did not Kármán gait, but rather held station using swimming kinematics that more closely resembled undulation during freestream swimming. Even so, we did observe a greater tail amplitude in the KVS array, with implications for vorticity control. We also observed certain kinematics differing from freestream treatment due to the effect of cylinders on the motion of the fluid, though not between cylinder arrays: namely, the COM amplitude increased in the PVS array and the snout amplitude increased in both the KVS array and the PVS array by similar amounts. It is possible that vortices were too dissipated in our zone of interest for fish to properly Kármán gait, or alternatively that the organization and morphology of the KVS array did not make Kármán gaiting feasible. Trout also commonly performed large, cross-stream searching motions before accelerating through the cylinder array, suggesting that sensory inputs can influence higher-order behaviors.

## List of Abbreviations and Symbols

KVS array A: cylinder array treatment that produced a Kármán vortex street
PVS array A: cylinder array treatment that produced a parallel vortex street
TKE: Turbulence kinetic energy
CFD: Computational Fluid Dynamics
COM: Center of mass
St: Strouhal number

## ACKNOWLEDGEMENTS

We would like to thank J. Strother for helpful discussions about this project and B. Ray for guidance with fish care.

## AUTHOR CONTRIBUTIONS

Conceptualization: A.C., J.C.L.; Methodology: D.S., E.R., A.C., J.C.L.; Validation: D.S., E. R., A.C., J.C.L.; Formal analysis: D.S., E.R.; Investigation: D.S., E.R., J.C.L; Resources: A.C., J.C.L.; Data curation: D.S., E.R.; Writing – original draft: D.S. E.R.; Writing – review & editing: D.S., E.R., A.C., J.C.L.; Visualization: D.S., E.R., A.C., J.C.L.; Supervision: A.C., J.C.L.; Project administration: J.C.L.; Funding acquisition: A.C., J.C.L.

## REFERENCES

1. Adams, B., Bohnhoff, W., Dalbey, K., Ebeida, M., Eddy, J., Eldred, M., Hooper, R., Hough, P., Hu, K., Jakeman, J., et al. (2020). Dakota, A Multilevel Parallel Object-Oriented Framework for Design Optimization, Parameter Estimation, Uncertainty Quantification, and Sensitivity Analysis: Version 6.13 User’s Manual. Albuquerque: Sandia National Lab (SNL-NM). doi:10.2172/1817318.

2. Akanyeti, O. and Liao, J. C. (2013a). The effect of flow speed and body size on Kármán gait kinematics in rainbow trout. J. Exp. Biol. 216, 3442–3449.

3. Akanyeti, O. and Liao, J. C. (2013b). A kinematic model of Kármán gaiting in rainbow trout. J. Exp. Biol. 216, 4666–4677.

4. Altringham, J. D. and Ellerby, D. J. (1999). Fish swimming: patterns in muscle function. J. Exp. Biol. 202, 3397–3403.

5. Bainbridge, R. (1962). Caudal fin and body movement in the propulsion of some fish. J. Exp. Biol. 40, 23–56.

6. Bak-Coleman, J., Court, A., Paley, D. A. and Coombs, S. (2013). The spatiotemporal dynamics of rheotactic behavior depends on flow speed and available sensory information. J. Exp. Biol. 216, 4011–4024.

7. Beal, D. N., Hover, F. S., Triantafyllou, M. S., Liao, J. C. and Lauder, G. V. (2006). Passive propulsion in vortex wakes. J. Fluid Mech. 549, 385–402.

8. Castro-Santos, T., Cotel, A. and Webb, P. W. (2009). Fishway evaluations for better bioengineering – an integrative approach. In Challenges for Diadromous Fishes in a Dynamic Global Environment, American Fisheries Society, Symposium. 69, pp. 557–575.

9. Di Santo, V., Goerig, E., Wainwright, D. K., Akanyeti, O., Liao, J. C., Castro-Santos, T. and Lauder, G. V. (2021). Convergence of undulatory swimming kinematics across a diversity of fishes. Proc. Natl. Acad. Sci. USA. 118, e2113206118.

10. Elder, J. and Coombs, S. (2015). The influence of turbulence on the sensory basis of rheotaxis. J. Comp. Physiol. A. 201, 667–680.

11. Enders, E. C., Boisclair, D. and Roy, A. G. (2003). The effect of turbulence on the cost of swimming for juvenile Atlantic salmon (*Salmo salar*). Can. J. Fish. Aquat. Sci. 60, 1149–1160.

12. Gao, Y., Chen, W., Wang, B. and Wang, L. (2020). Numerical simulation of the flow past six-circular cylinders in rectangular configurations. J. Mar. Sci. Technol. 25, 718–742.

13. Gillis, G. B. (1998). Neuromuscular control of anguilliform locomotion: patterns of red and white muscle activity during swimming in the American eel *Anguilla rostrata*. J. Exp. Biol. 201, 3245–3256.

14. Hammond, L., Altringham, J. D. and Wardle, C. S. (1998). Myotomal slow muscle function of rainbow trout *Oncorhynchus mykiss* during steady swimming. J. Exp. Biol. 201, 1659–1671.

15. Igarashi, T. (1981). Characteristics of the flow around two circular cylinders arranged in tandem: 1st report. Bull. JSME. 24, 323–331.

16. Jayne, B. C. and Lauder, G. V. (1995). Are muscle fibers within fish myotomes activated synchronously? Patterns of recruitment within deep myomeric musculature during swimming in largemouth bass. J. Exp. Biol. 198, 805–815.

17. Karasudani, T. and Funakoshi, M. (1994). Evolution of a vortex street in the far wake of a cylinder. Fluid Dyn. Res. 14, 331.

18. Kassambara, A. (2021). rstatix: Pipe-friendly framework for basic statistical tests. R package version 0.7.0. https://CRAN.R-project.org/package=rstatix.

19. Kennedy, J. S. (1983). Zigzagging and casting as a programmed response to wind-borne odour: a review. Physiol Entomol. 8, 109–120.

20. Lacey, R. W. J., Neary, V. S., Liao, J. C., Enders, E. C. and Tritico, H. M. (2012). The IPOS framework: linking fish swimming performance in altered flows from laboratory experiments to rivers. River Res. Appl. 28, 429–443.

21. Lauder, G. V. and Drucker, E. G. (2004). Morphology and experimental hydrodynamics of fish fin control surfaces. IEEE J. Ocean Eng. 29, 556–571.

22. Liao, J. C. (2004). Neuromuscular control of trout swimming in a vortex street: implications for energy economy during the Kármán gait. J. Exp. Biol. 207, 3495– 3506.

23. Liao, J. C. (2006). The role of the lateral line and vision on body kinematics and hydrodynamic preference of rainbow trout in turbulent flow. J. Exp. Biol. 209, 4077–4090.

24. Liao, J. C. (2007). A review of fish swimming mechanics and behaviour in altered flows. Philos. Trans. R. Soc. B. 362, 1973–1993.

25. Liao, J.C. (2022). Fish swimming efficiency. Curr. Biol. 32, R589–R683.

26. Liao, J. C., Beal, D. N., Lauder, G. V. and Triantafyllou, M. S. (2003a). The Kármán gait: novel body kinematics of rainbow trout swimming in a vortex street. J. Exp. Biol. 206, 1059–1073.

27. Liao, J. C., Beal, D. N., Lauder, G. V. and Triantafyllou, M. S. (2003b). Fish exploiting vortices decrease muscle activity. Sci. 302, 1566–1569.

28. Lucas, K. N., Lauder, G. V. and Tytell, E. D. (2020). Airfoil-like mechanics generate thrust on the anterior body of swimming fishes. Proc. Natl. Acad. Sci. USA. 117, 10585–10592.

29. Mathis, A., Mamidanna, P., Cury, K. M., Abe, T., Murthy, V. N., Mathis, M. W. and Bethge, M. (2018). DeepLabCut: markerless pose estimation of user-defined body parts with deep learning. Nat. Neurosci. 21, 1281–1289.

30. Nath, T., Mathis, A., Chen, A. C., Patel, A., Bethge, M. and Mathis, M. W. (2019). Using DeepLabCut for 3D markerless pose estimation across species and behaviors. Nat. Protoc. 14, 2152–2176.

31. Przybilla, A., Kunze, S., Rudert, A., Bleckmann, H. and Brücker, C. (2010). Entraining in trout: a behavioural and hydrodynamic analysis. J. Exp. Biol. 213, 2976–2986.

32. Puzdrowska, M. and Heese, T. (2019). Turbulent kinetic energy in bolt fishway. AgriEngineering. 1, 265–282.

33. Stewart, W. J., Tian, F., Akanyeti, O., Walker, C. J. and Liao, J. C. (2016). Refuging rainbow trout selectively exploit flows behind tandem cylinders. J. Exp. Biol. 219, 2182–2191.

34. Taguchi, M. and Liao, J. C. (2011). Rainbow trout consume less oxygen in turbulence: the energetics of swimming behaviors at different speeds. J. Exp. Biol. 214, 1428– 1436.

35. Triantafyllou, M. S., Triantafyllou, G. S. and Yue, D. K. P. (2000). Hydrodynamics of fishlike swimming. Annu. Rev. Fluid Mech. 32, 33–53.

36. Tritico, H. M. and Cotel, A. J. (2010). The effects of turbulent eddies on the stability and critical swimming speed of creek chub (*Semotilus atromaculatus*). J. Exp. Biol. 213, 2284–2293.

37. Wardle, C., Videler, J. and Altringham, J. (1995). Tuning in to fish swimming waves: body form, swimming mode and muscle function. J. Exp. Biol. 198, 1629–1636.

38. Webb, P. W. (2002). Control of posture, depth, and swimming trajectories of fishes. J. Exp. Biol. 42, 94–101.

39. Webb, P. W. (1988). ‘Steady’ swimming kinematics of tiger musky, an esociform accelerator, and rainbow trout, a generalist cruiser. J. Exp. Biol. 138, 51–69.

40. Webb, P. W. (1998) Entrainment by river chub *Nocomis micropogon* and smallmouth bass *Micropterus dolomieu* on cylinders. J. Exp. Biol. 201, 2403–2412.

41. Webb, P. W. and Weihs, D. (1994). Hydrostatic stability of fish with swim bladders: not all fish are unstable. Can. J. Zool. 72, 1149–1154.

42. Wilkes, M. A., Enders, E. C., Silva, A. T., Acreman, M. and Maddock, I. (2017). Position choice and swimming costs of juvenile Atlantic salmon *Salmo salar* in turbulent flow. J. Ecohydraulics. 2, 16–27.

43. Wu, T. Y. and Chwang, A. T. (1975). Extraction of flow energy by fish and birds in a wavy stream. In Swimming and Flying in Nature: Volume 2 (ed. T. Y.-T. Wu, C. J. Brokaw and C. Brennen), pp. 687–702. Boston, MA: Springer US.

44. Xiong, G. and Lauder, G. V. (2014). Center of mass motion in swimming fish: effects of speed and locomotor mode during undulatory propulsion. Zoology. 117, 269–281.

